# The genomic basis of electrotaxis in *Dictyostelium discoideum*: Electric field sensitive amino acids are dynamically encoded en masse for the streaming-stage proteome

**DOI:** 10.1101/034892

**Authors:** Albert J. Erives

## Abstract

Electrotaxis plays a critical role in developmental cell migration, axon growth cone guidance, epithelial wound healing, tissue regeneration, and the degree of invasiveness characterizing different cancer cell lines. During electrotaxis in a direct current electric field (EF), a cell migrates preferentially either towards the anode or cathode depending on the cell-type. However, the types and ranges of mechanisms coupling trans-cellular electric fields to cellular EF-sensitive signaling systems are largely unknown. To address this cell biological phenomenon, I use transcriptomic data from a developmental genetic model in which multicellular social aggregation is induced by starvation of amoeboid cells. I find that the developmental proteome expressed during the streaming aggregation stage is measurably and substantially enriched in charged and highly polar amino acids relative to the proteomes of either the unicellular amoeboid or the multicellular fruiting body. This large-scale coding augmentation of EF-sensitive amino acid residues in thousands of streaming-specific proteins is accompanied by a proportional coding decrease in the number of small, uncharged amino acid residues. I also confirm an expected coding increase of biosynthetically costly amino acids in the proteome of the satiated feeding-stage amoeboid. These findings suggest that electrotactic capability is encoded broadly in the genetically regulated deployment of a developmental proteome with augmented EF-sensitivity. These results signify that extreme, nonuniform, evolutionary constraints can be exerted on the amino acid composition of an organism’s proteome.

## Introduction

Electrotaxis, also known as galvanotaxis, is a key aspect of developmental cell migration, growth cone axon guidance, epithelial wound healing and/or tissue regeneration, and the degree of invasiveness characterizing different cancer cell lines (Jaffe and Nuccitelli 1977; Nuccitelli *et al*. 1977; McCaig *et al*. 2005; Shanley *et al*. 2006; Tsai *et al*. 2013). Depending on the cell type, during electrotaxis a cell migrates towards the cathode or anode defining a direct current electric field (EF). Many well-studied examples of electrotaxis were discovered initially with the experimental application of exogenous electric current. However, endogenous electric fields have been identified as well. In most cases, such as the social aggregation of *Dictyostelium* cells or *Xenopus* tail regeneration, endogenous EFs are established in a regulated fashion through asymmetric membrane depolarization in individual cells thus leading to directional ion current flow across a field of cells (Anderson 1962; Nuccitelli *et al*. 1977; McCaig *et al*. 2005; Reid *et al*. 2009; Zhang *et al*. 2015). Nonetheless, as important as electrotaxis is to understanding the basic biology of cells, the range and dimensions of its underlying mechanisms and the existence of possible unifying principles coupling external electric fields to cellular EF-sensing and signaling systems remain largely unknown.

Social slime molds are ideal model developmental genetic systems to study the basic principles underlying electrotaxis (Fig. 1). Diverse social slime molds, including *Dictyostelium* and *Physarum* species, exhibit electrotaxis towards the cathode direction when streaming into a multicellular aggregation or as a syncytium, respectively, eventually forming a fruiting body (Anderson 1962; Nuccitelli *et al*. 1977). Numerous studies have documented the asymmetric distribution of both common cations and specific proteins in either the leading edge or trailing cellular processes (Anderson 1962; Coates and Harwood 2001). Furthermore, more recent studies have also begun to disentangle the specific proteins involved in electrotaxis versus chemotaxis (Coates and Harwood 2001; Zhao *et al*. 2002; Shanley *et al*. 2006; Myre *et al*. 2011; Wessels *et al*. 2014). More recently, systematic genetic screens for genes underlying electrotaxis have also been conducted (Gao *et al*. 2015). However, in principle, diverse cellular proteins involved in cell motility, directional orientation, regulation of membrane current flow, and other signaling systems will likely evolve to be oriented or subcellularly localized in an optimal fixed relationship to the direction of electrotaxis and/or cell polarity (Fig. 1B). Because proteins can evolve to have different complements and distributions of charged and polarized amino acid residues, evolution is likely to achieve optimal protein EF preferences in part by the selection of amino acid composition in addition to active transport and active capture mechanisms for each protein that needs to be asymmetrically localized or oriented. This expectation of skewed amino acid composition towards EF-sensitivity is in addition to the known role of charged residues in voltage-gated membrane channels, such as the four arginine (Arg, R) residues in Kv1.2 potassium channels, which transduce changes in membrane potential (Zhang *et al*. 2015). Thus, the specific question, which I ask and address in this report, is to determine the extent to which this macroscopic proteomic principle for EF-sensitivity is manifested and detectable in a developmental genetic model of electrotaxis.

**Figure 1.**
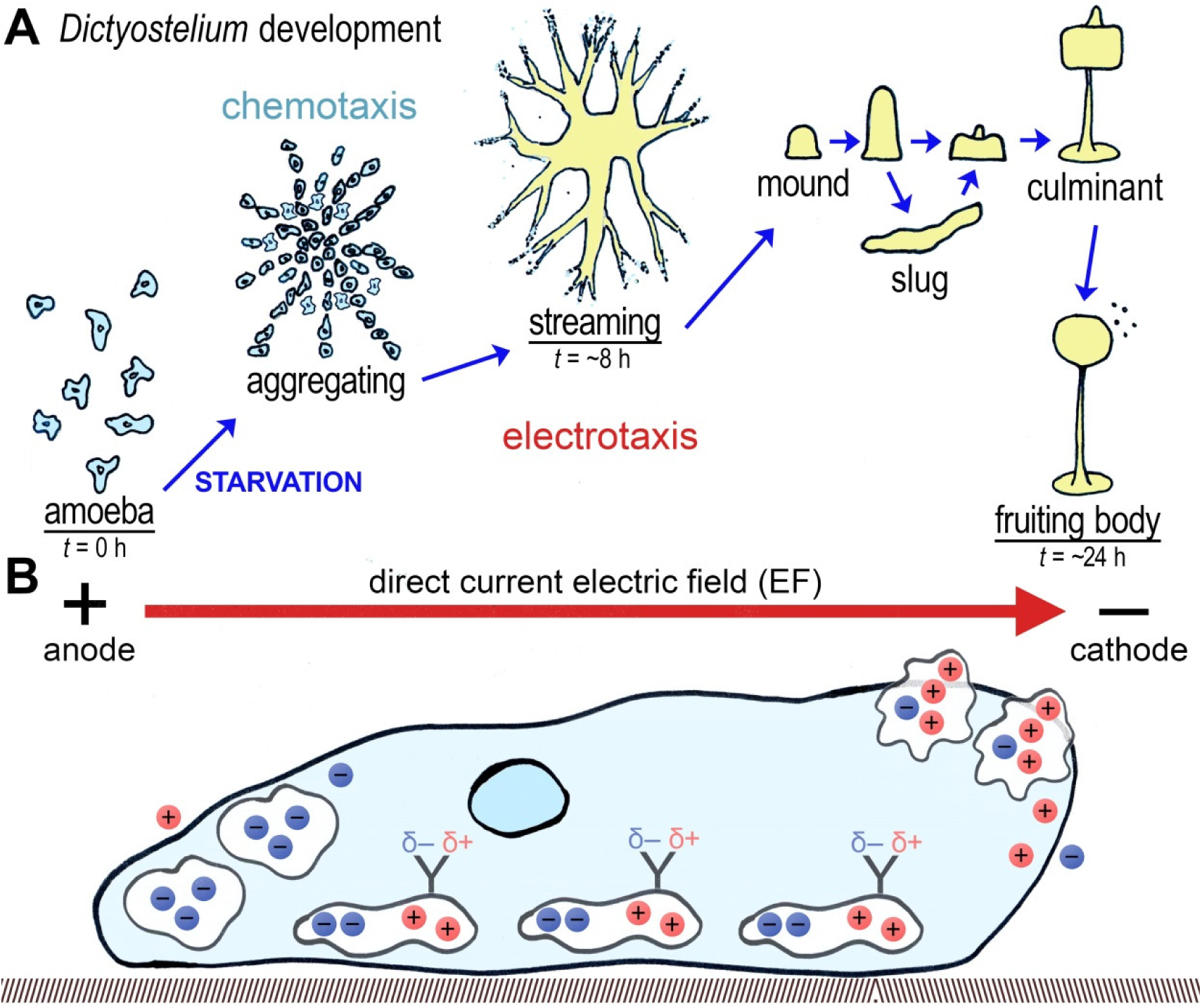
Electrotaxis during *Dictyostelium* development. (**A**) Depicted are the developmental stages of *Dictyostelium discoideum*, a slime mold and model genetic system. Development is induced by starvation of the amoeboid form, upon which amoeboid cells aggregate via chemotaxis. Amoeboid cells are also sensitive to direct current electric fields (EF) and will spontaneously stream in the direction of the cathode, similar to the natural process of developmental streaming. Streaming cells coalesce into a mound that develops into the fruiting body, which eventually releases spores. The stage-specific proteomes for the amoeba, streaming aggregate and the fruiting body (underlined) are analyzed in this study. (**B**) Depicted is a single *Dictyostelium* cell migrating by electrotaxis towards the cathode and three types of intracellular and/or membrane-bound proteins that differ in their net complement of fixed positive (red) and negative (blue) charges from their constituent amino acid residues. Proteins with net positive charges will migrate electrophoretically towards the leading edge of the cell. This process is postulated to lead to the asymmetric distribution of key membrane proteins required for environmental sensing and cell motility. Proteins with net negative charges would migrate electrophoretically towards the rear of the cell. Last, proteins with net neutral charges but asymmetric distributions of charged residues would align themselves with the electric field lines accordingly, as would the sidechains of uncharged polar amino acids (*e.g*., asparagine and glutamine depicted by “Y” shaped end groups and asymmetric charge distributions).

Here I use transcriptomic data sets from the developmental stages of *Dictyostelium discoideum*, whose multicellular social aggregations are induced by starvation experienced in the solitary amoeboid forms. I find that the proteome-specific to the streaming stages is significantly enriched in all EF-sensitive amino acids (charged and polar uncharged amino acids) relative to solitary amoeboid-specific and fruiting body-specific proteomes. This increase in the proportional coding of these amino acids in the streaming-specific proteins is achieved by a corresponding decrease in the coding of small, uncharged amino acids. These observations are also supported by confirmation of a second expected genomic signature: decreased coding of biosynthetically costly amino acids during the multicellular developmental stages relative to the solitary amoeboid form, whose stage-specific proteome is deployed under non-starvation conditions. These findings suggest that electrotaxis is diffusively encoded in the genetically regulated expression of diverse proteins with measurable increases in the proportion of EF-sensitive amino acid side-chains relative to other developmental stages.

## Material and Methods

Transcriptomic analyses. The NCBI GEO dataset GSE71036 (Gokhale, R.S. *et al*.), which includes developmental-stage transcriptomic profiling of wild type *Dictyostelium discoideum* (AX4 strain) was analyzed using GEO2R (Barrett *et al*. 2011; Barrett *et al*. 2013). First, distributions of the signal values for two replicates of each of three stages (amoeboid, streaming, and fruiting body stages) were plotted to determine and confirm that the data set had normalized medians and are appropriately cross-comparable (Fig. S1). Then GEO2R was used to identify data sets for all pair-wise comparisons of the three developmental groups. A Benjamini & Hochberg (False discovery rate) adjustment of P-values was applied to focus on stage-specific genes with values ≤ 0.05. Similar procedure was used in the analysis of GSE42407 and GSE62146, except one of three replicate samples from each cell line was dropped in GSE42407 because their median distributions were discordant from the majority of samples. Figure S2 shows the distributions of the samples actually used.

**Amino acid compositional analysis**. The BioMart database query system for *protists.ensembl.org* was then used to retrieve those genes encoding a unique peptide sequence (Haider *et al*. 2009; Jones *et al*. 2011; Smedley *et al*. 2015). Using UNIX command lines as well as perl and vim, FASTA headers, ambiguous characters (“X” and “U”), and a sporadic number of FASTA headers with the sequence “Sequence unavailable” were removed. All 20 amino acid characters while replacing each character with lower-case versions. After counting the characters, stop codon and newline characters (\*, \n) were removed to count the total number of residues in each data set. This total sum was then used to doublecheck the sum of the individual amino acid character counts.

**Distance-based dendrogram of amino acid composition**. Percentages of each of the three amino acid groups depicted in Fig. 2A were used to build a distance matrix, which was then used to build a tree by Fitch-Margoliash algorithm using PhyloDraw software (Choi *et al*. 2000).

**Figure 2.**
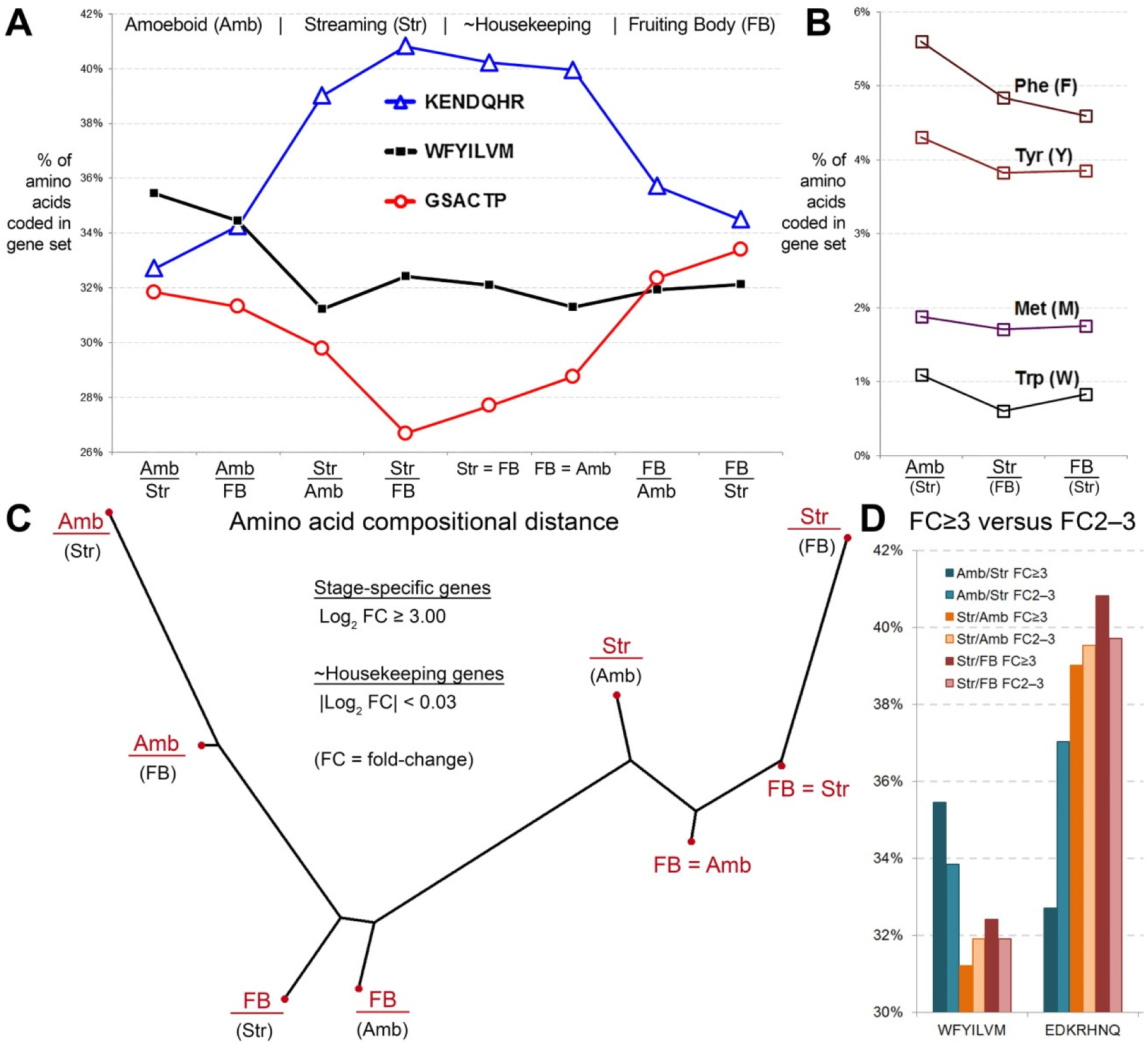
Substantial differences in amino acid coding frequencies distinguish the different developmental stages of *Dictyostelium*. (**A**) Shown are the percentage of three groups of amino acids in stage-specific and house-keeping genes. Stage specific genes were identified as those expressed more than eight-fold higher relative to another stage (log_2_ fold-change ≥ 3.00). Housekeeping genes represent genes expressed at similar levels in two stages (log_2_ fold-change < 0.03). See material and methods for number of genes in each group. Graph shows that the amoeboid cells are less constrained in their usage of biosynthetically-expensive amino acids (black line), while streaming stages are enriched in electro-sensitive amino acids (blue line) at the expense of uncharged small amino acids (red line). (**B**) Graph of individual biosynthetically expensive amino acids by the three main stages. All lines are color-coded and listed in the keys in order from top to bottom as they appear on the graphs. (**C**) A pairwise distance matrix based on amino acid composition levels shown in (A) was calculated for all eight proteomic sets and used to construct a dendrogram using the Fitch-Margoliash algorithm. This distance-based tree demonstrates the developmental coherency of amino acid composition and the unique change in the streaming stage proteome. (D) Protein-coding genes expressed at log FC 2–3 recapitulate amino acid composition for genes expressed at log FC ≥ 3. Shown are biosynthetically expensive amino acids (WFYILVM) on the left, and EF-sensitive amino acids (EDKRHNQ) on the right. The log FC 2–3 genes are shown in a lighter shade than the log FC ≥ 3 genes.

## Results and Discussion

### A model genetic system to understand systemic developmental regulation of electrotaxis

Here I describe novel, striking, and straightforward findings concerning a basic principle of electrotaxis by cells and the manner for its evolution. These findings also suggest general considerations and approaches to identifying the extent of electric-field sensitive proteomes in diverse developmental-stage specific proteomes. I arrived at these findings after hypothesizing that the protein-coding genes differentially expressed during development of the multi-fruiting body of a social slime mold (Fig. 1A) should show some degree of differential constraints of amino acid codon usage for two simple reasons. First, chemotaxis and loose aggregation in the solitary feeding amoeba form is induced by starvation. Thus, proteins differentially expressed specifically in the satiated vegetative amoeboid state should be the least constrained in their usage of biosynthetically costly amino acids relative to the proteins of the social, multicellular, developmental stages.

Second, I postulated that the inherent behavioral potential for electrotactic motility of vegetative amoeba in direct current electric fields (EF) implies that it is used dynamically in the life cycle. This capability is likely most active during streaming aggregation, during which cells become physically coupled by polarized adherens junctions (Coates and Harwood 2001) and the cells are at their peak motility and capable of being directed by EFs. This second constraint is also indirectly implied by current hypotheses of receptor allocation during electrotaxis in amoeba, migrating mammalian epithelial cells, and growth cone taxis (McCaig *et al*. 2005), and is supported by studies of electrotaxis in various aggregating slime molds (Jaffe and Nuccitelli 1977; Nuccitelli *et al*. 1977). Thus, electrotactic cells likely benefit from intrinsic electrophoretic capability for diverse cathodal and anodal directed proteins (Fig. 1B, charged proteins at front or back of polarized cells). Electrotactic cells also likely benefit from intrinsic polarization of neutral proteins that are optimally oriented relative to EF current lines (also shown in Fig. B). Last, many membrane-spanning proteins involved in electrotaxis, such as voltage-gated channels or receptors of highly charged small molecules, will require intrinsically charged amino acid residues (Nakajima *et al*. 2015; Zhang *et al*. 2015). Altogether, there are many reasons for expecting intrinsic EF-sensitivity for diverse proteins expressed during electrotaxis.

To test for differential biosynthetic and electrotactic constraints, I considered the frequencies of amino acids from three major groups. The first group consisted of the hydrophobic amino acids, which includes most of biosynthetically costly amino acids: Trp (W), Phe (F), Tyr (Y), Ile (I), Leu (L), Val (V), and Met (M). The second group included all of the EF-sensitive amino acids (*i.e*., the charged amino acids and the more polar uncharged amino acids): Lys (K, +), Arg (R, +), His (H, [+]) Glu (E, −), Asp (D, −), Gln (Q, +/−), and Asn (N, +/−). The third group consisted of mostly the small uncharged amino acids: Gly (G), Ser (S), Thr (T), Ala (A), Cys (C), and Pro (P).

To determine the extent to which there are differential constraints on amino acid usage by developmental stage, I analyzed gene expression data for three distinct developmental stages of wild type *Dictyostelium discoideum* (AX4): Amoeboid (“Amb”, *t* = 0 hours post-starvation), Streaming (“Str”, t = ~8 hours post-starvation), and Fruiting Body (“FB”, *t* = 24 hours post-starvation). For all such stages, I identified the set of protein-coding genes that are expressed at eight-fold higher levels relative to levels at another stage (log _2_ fold-change ≥ 3.00). These sets ranged from 83 protein-coding genes up to 680 protein-coding genes (see Table 1). I also looked at 411 genes expressed at nearly the same levels in streaming and fruiting body stages (254,342 residues), and 327 genes expressed at nearly the same levels in amoeboid and fruiting body stages (188,416 residues).

**Table 1.** Stage-specific and multi-stage constitutive gene sets analyzed. Gene sets were identified by identifying transcripts expressed at eight-fold higher levels in one stage relative to another [log fold-change (FC) ≥ 3.00 (a), or ≥ 2 but < 3 (b)] or by those expressed at similar levels in two different stages *(e.g*., log FC FB/Amb < 0.03). The number of genes and their total number of encoded amino acid residues are indicated for each gene set. In some figures, the constitutive gene sets are labeled as “-Housekeeping”.

| Gene Set* | Number of protein-coding genes | Number of encoded amino acid residues |
| --- | --- | --- |
| Amoeboid 1a (log FC over Str $\geq 3$) | 108 | 45,188 |
| Amoeboid 1b (log FC over Str 2–3) | 422 | 209,922 |
| Amoeboid 2 (log FC over FB $\geq 3$) | 83 | 30,637 |
| Streaming 1a (log FC over Amb $\geq 3$) | 398 | 157,981 |
| Streaming 1b (log FC over Amb 2–3) | 323 | 159,678 |
| Streaming 2a (log FC over FB $\geq 3$) | 140 | 49,488 |
| Streaming 2b (log FC over FB 2–3) | 413 | 182,074 |
| Constitutive 1 (log FC FB/Str $< 0.03$) | 411 | 254,342 |
| Constitutive 2 (log FC FB/Amb $< 0.03$) | 327 | 192,515 |
| Fruiting Body (log FC over Amb $\geq 3$) | 680 | 225,087 |
| Fruiting Body (log FC over Str $\geq 3$) | 418 | 127,202 |

### Constraint on biosynthetically expensive amino acids in the stages induced by starvation

I found that the protein-coding transcriptome specifically expressed in the amoeboid form is the least constrained in its usage of biosynthetically costly amino acids. This amino acid group is encoded at 36% of all amino acids in the amoeboid stage while it is decreased to ~32% in streaming and fruiting body specific proteomes and in non-stage specific proteomes (Fig. 2A). This can be broken down into the most costly amino acids of Phe, Tyr, Met, and Trp, all of which show higher levels in the amoeboid stage (Fig. 2B). Thus, this result confirms the prediction of relaxed selective constraints for biosynthetically expensive amino acids in the feeding (i.e., satiated) state of the solitary amoeboid form of a facultative social slime mold.

### Only EF-sensitive amino acids are enriched in the streaming stage proteome

A much more substantial change was observed for the EF-sensitive amino acids, including the charged amino acids (E−, D−, K+, R+, H[+]) and the most polar uncharged amino acids (N, Q). I found that the EF-sensitive amino acid set was encoded at the highest levels in the differentially expressed proteomes of the streaming stage regardless of whether the comparison is to amoeboid or fruiting body stages (Fig. 2A). There is an increase of +7.1% in the EF-sensing amino acid set from the amoeboid stage (32.7%) to its peak levels in the streaming stage (40.8%), representing a staggering 21.7% proportional increase (7.1% of 32.7%). This finding is consistent with the above proposed idea and previous results that electrotaxis is optimized for the streaming stages. I also found streaming-stage enrichment for each EF-sensitive amino acid (Fig. 3A). If these are sub-grouped separately into the uncharged polar (N, Q), the positively charged (K, R, [H]), and negatively charged (E, D) amino acids, each subset is also found to have identical developmental coding profiles (Fig. 3B). This suggests that the EF-sensitivity of charged and polar side-chains is the important common property of the cellular proteome during electrotaxis. This also suggests that electrotaxis is genetically wired diffusely throughout the genome in both gene regulatory sequences and a protein-coding repertoire enriched in codons for charged and polar uncharged amino acids. Because this strong signature is also seen in the protein-genes that are expressed at the same levels in different stages (Fig. 2A), it implies that the selection for this generalized enrichment of EF-sensitive amino acid residues is only ever relaxed in proteins that are not expressed in the streaming stages.

**Figure 3.**
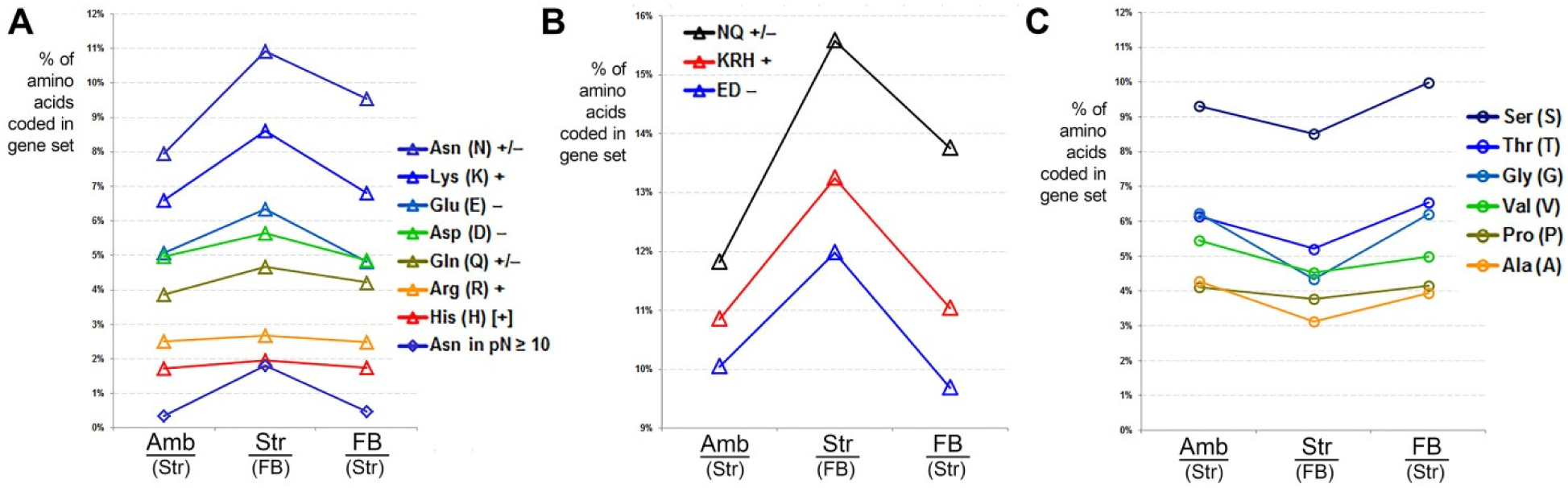
Frequency of all electric field-sensitive amino acids are increased in the streaming-specific proteome. (**A**) Individual charged and uncharged polar amino acids are shown. The percentage of all amino acids is highest in streaming cells. Interestingly, most of the increase in Asp (N) is the result of polyN tracts (see text for discussion of import). (**B**) A breakdown of the EF-sensitive amino acids by charge type: polar uncharged (N, Q), positively charged (K, R, [H]), and negatively charged (E, D). Usage of each group is increased at similar levels at the coding level. (**C**) The increase in EF-sensitive amino acids depicted in A and B are evidently offset by a decrease in small uncharged amino acids (S, T, G, V, P, A).

I found that the enrichment of the EF-sensitive amino acids comes at the expense of the small uncharged amino acids, which exhibit the opposite trend as the EF-sensitive amino acids (Fig. 3C). This suggests that most of these amino acids (except perhaps C and S) are more dispensable than the large aliphatic and aromatic amino acids of the biosynthetically costly group. Alternatively, these amino acids can be referred to as “EF-refractory” and their enrichment at specific stages would in general diminish the capacity of a cell to be externally manipulated by local electric fields.

To quantify the apparent temporal coherency in the dynamics of amino acid composition during *Dictyostelium* development, I also constructed a distance matrix for all pairwise comparisons of the analyzed data sets using the levels of the three amino acid groups: (i) biosynthetically expensive aliphatic and aromatic amino acids, (*ii*) the EF-sensitive amino acids, and (*iii*) the EF-refractory amino acids. I then used a weighted least-squares method (Fitch-Margoliash) to cluster the eight proteomes into a dendrogram of related amino acid compositions (Fig. 2C). This tree indeed shows that similar data sets from the same developmental time points are typically more like each other in amino acid composition than they are to other time points. Thus, developmental change in EF-sensitive amino acid composition is both measurable and dynamically coherent. I thus propose that this regulated switch underlies a gradual change from a strongly chemotactic but weakly electrotactic amoeboid state to a strongly electrotactic streaming state. Certainly, specific key proteins will be involved in the localized control of trans-membrane ionic currents and membrane potential. Nonetheless, the proposed hypothesis for an EF-sensitive proteome of electrotaxis implies that evolutionary selective pressure sculpts the amino acid composition of thousands of protein-coding genes expressed at a developmental stage.

To confirm the proteome-wide EF-sensitivity hypothesis for *Dictyostelium* development, I then analyzed the amino acid composition for genes expressed at the same stages but with a more modest log fold-change between two and three (4x–8x higher than another stage; see Table 1). I found that these new protein-coding gene sets recapitulated the salient features of the most stage-specific genes (log FC ≥ 3): (*i*) biosynthetically expensive amino acids are used more freely in the amoeboid stages, and (*ii*) EF-sensitive amino acids are enriched in the streaming aggregation stages (Fig. 2D). Thus, the results obtained with the most stage-specific genes are not an artifact of a having chosen a random set of developmental genes.

### The polyN but not the polyQ proteome is developmentally enriched in the streaming stage

I also found that most of the streaming-stage increase in asparagine (N) is due to residues in contiguous polyN tracts that are ≥ 10 residues long (Fig. 3A). Adaptive proteostatic systems have been postulated to attenuate aggregation of the extensive polyN repertoire in *Dictyostelium* (Malinovska *et al*. 2015). PolyN tracts, like polyQ tracts have been implicated in protein aggregation and are often potent prionizing domains (Peters and Huang 2007). However, unlike glutamine, asparagine has a reduced propensity to form protein secondary structure (α-helices and β-sheets). Furthermore, the precise lengths of polyQ tracts in specific signaling factors such as the Notch intracellular domain that functions in transcriptional co-activation may be more important in specific protein-protein interactions than polyN tracts (Rice *et al*. 2015). Thus, polyQ tracts may evolve by additional constraints not applicable to polyN tracts.

To explore further the apparent streaming-stage specific enrichment for polyN content, I also plotted polyQ and polyN content for all of the developmental genes sets (Fig. 4). This developmental profile shows that polyN proteome is largely streaming stage-specific. This is not observed for polyQ tracts, which are generally and modestly enriched only in proteomic sets that are expressed at similar levels at different stages. This strongly suggests either a functional or a physiological significance to increasing bulk polyN content during the streaming stage. Thus, it will be interesting to determine in future studies whether there are more specific roles for polyN tracts in electrotaxis than simply featuring electrically polarized side-chains. Interestingly, it has been previously found that the highly conserved, polyN and polyQ-containing Huntingtin protein of *Dictyostelium*, whose homolog in humans is involved in protein aggregation and neurodegeneration, is required for streaming aggregation (Myre *et al*. 2011; Wessels *et al*. 2014).

**Figure 4.**
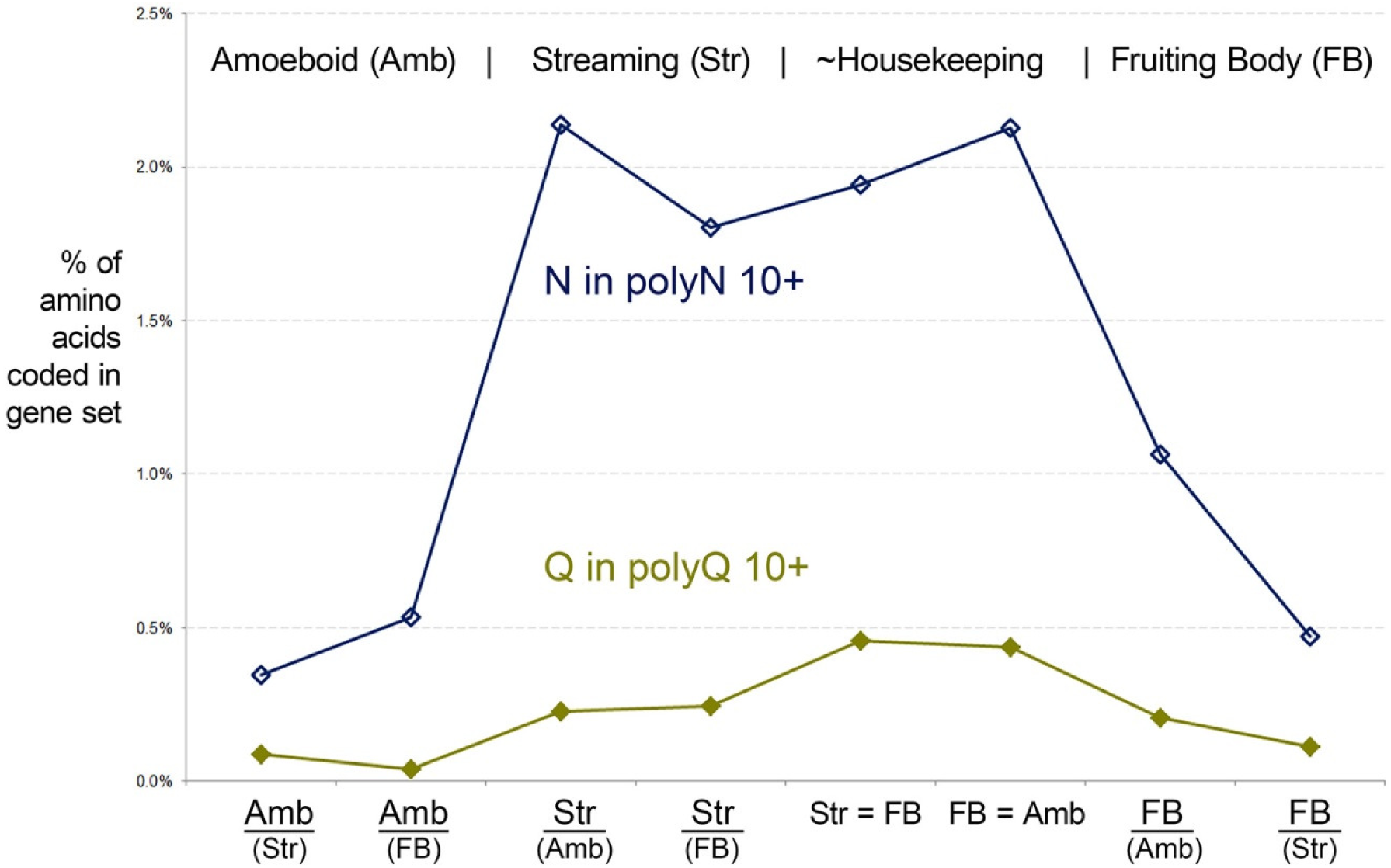
The polyN proteome but not the polyQ proteome is largely specific to streaming. Shown is the percentage of all amino acids that are N or Q and exist in polyN (dark blue line) or polyQ (olive line) tracts that are 10 residues or longer. Future studies will need to determine the biological significance of polyN-rich proteins in *Dictyostelium* biology.

### Developmental regulation of proteomic EF-sensitivity in *Dictyostelium*, may be relatively extreme

The described findings may serve as a general reference guide for disentangling the various evolutionary constraints impinging on the broad EF-sensitivity of mammalian cells during developmental cell migration or pathological metastasis. One might expect this to be the case given that current studies are identifying independent and parallel pathways by which EFs polarize even a single cell type under different conditions (Saltukoglu *et al*. 2015). Alternatively, it is possible that natural selection has grossly and atypically exaggerated EF-sensitivity in the streaming-stage proteome of *Dictyostelium*. To begin to assess this question, I conducted equivalent compositional analyses for two additional transcriptomic series. The first transcriptomic data set (GSE42407) comes from a series of human lung cancer cell lines that were derived from a parent line, CL1-0, by five rounds of laboratory selection for increasing matrix invasion culminating in an aggressive CL1-5 line (Chen *et al*. 2013). CL1-5 has been found to be anodally electrotactic unlike the less-aggressive parent CL-0. In a second transcriptomic data set (GSE62146), I compared mouse iPS fibroblast cells to those induced to form neurons via over-expression of *NEUROGENIN1*.

Results for both mammalian data sets show that changes in amino acid composition are more modest and mixed than those seen during *Dictyostelium* development (Table 2). Possibly, these data sets compare cell lines that are too similar in developmental profile and phenotype. But in any case, these analyses cannot rule out whether proteome-wide EF-sensitive augmentation yet exists in specific developmental contexts in some animals, or whether the electrotactic proteome of *Dictyostelium* is an idealized extreme manifestation.

**Table 2.** Developmental regulation of proteomic EF-sensitivity may be augmented to exceptional levels in *Dictyostelium*. The top over and under represented amino acids in dataset B relative to dataset A are highlighted in blue and red, respectively. See text for details.

| amino acids | 488 Amb genes<br>logFC≥3 | 398 Str genes<br>logFC≥3 | \|A-B\| | log(A/B) | 428 CL1-0 genes<br>logFC≥3 | 707 CL1-5 genes<br>logFC≥3 | \|A-B\| | log(A/B) | 1332 iPS genes<br>logFC≥3 | 1345 iPS+NGN1 genes<br>logFC≥3 | \|A-B\| | log(A/B) |
| --- | --- | --- | --- | --- | --- | --- | --- | --- | --- | --- | --- | --- |
| A | 4.27% | 2.90% | 1.37% | -0.56 | 6.88% | 6.78% | 0.10% | -0.02 | 6.96% | 7.38% | 0.42% | 0.09 |
| C | 1.75% | 2.09% | 0.34% | 0.26 | 2.21% | 2.28% | 0.06% | 0.04 | 2.30% | 2.22% | 0.08% | -0.05 |
| D | 4.97% | 4.89% | 0.08% | -0.02 | 4.99% | 4.73% | 0.27% | -0.08 | 4.73% | 4.90% | 0.17% | 0.05 |
| E | 5.07% | 5.41% | 0.34% | 0.09 | 6.82% | 7.15% | 0.33% | 0.07 | 6.92% | 7.10% | 0.17% | 0.04 |
| F | 5.60% | 4.71% | 0.89% | -0.25 | 3.61% | 3.52% | 0.09% | -0.04 | 3.48% | 3.44% | 0.04% | -0.02 |
| G | 6.24% | 4.65% | 1.59% | -0.42 | 6.65% | 6.93% | 0.28% | 0.06 | 6.86% | 6.84% | 0.02% | 0.00 |
| H | 1.73% | 1.84% | 0.11% | 0.09 | 2.55% | 2.58% | 0.03% | 0.02 | 2.45% | 2.58% | 0.13% | 0.07 |
| I | 8.60% | 8.35% | 0.25% | -0.04 | 4.47% | 4.34% | 0.13% | -0.04 | 4.19% | 4.22% | 0.03% | 0.01 |
| K | 6.60% | 7.21% | 0.61% | 0.13 | 5.61% | 5.65% | 0.04% | 0.01 | 5.63% | 5.36% | 0.28% | -0.07 |
| L | 8.51% | 7.94% | 0.57% | -0.10 | 9.27% | 9.70% | 0.43% | 0.07 | 9.94% | 9.64% | 0.30% | -0.04 |
| M | 1.88% | 1.51% | 0.37% | -0.32 | 2.23% | 2.12% | 0.11% | -0.07 | 1.96% | 2.08% | 0.12% | 0.09 |
| N | 7.96% | 12.56% | 4.60% | 0.66 | 3.94% | 3.80% | 0.14% | -0.05 | 3.53% | 3.56% | 0.03% | 0.01 |
| P | 4.12% | 4.22% | 0.10% | 0.03 | 6.56% | 6.43% | 0.12% | -0.03 | 6.62% | 6.60% | 0.02% | 0.00 |
| Q | 3.86% | 4.66% | 0.80% | 0.27 | 4.67% | 4.81% | 0.14% | 0.04 | 4.57% | 4.64% | 0.08% | 0.02 |
| R | 2.52% | 2.45% | 0.07% | -0.04 | 5.34% | 5.47% | 0.13% | 0.03 | 5.41% | 5.62% | 0.21% | 0.06 |
| S | 9.32% | 10.05% | 0.73% | 0.11 | 8.56% | 8.67% | 0.11% | 0.02 | 8.50% | 8.73% | 0.22% | 0.04 |
| T | 6.14% | 5.86% | 0.28% | -0.07 | 5.88% | 5.52% | 0.36% | -0.09 | 5.99% | 5.39% | 0.60% | -0.15 |
| V | 5.45% | 4.45% | 1.00% | -0.29 | 5.99% | 5.73% | 0.26% | -0.06 | 6.18% | 5.87% | 0.31% | -0.07 |
| W | 1.10% | 0.64% | 0.46% | -0.78 | 1.07% | 1.16% | 0.09% | 0.12 | 1.19% | 1.20% | 0.02% | 0.02 |
| Y | 4.31% | 3.61% | 0.70% | -0.26 | 2.68% | 2.62% | 0.06% | -0.03 | 2.57% | 2.62% | 0.05% | 0.03 |
| sum | 100% | 100% | 15.26% |  | 100% | 100% | 3.27% |  | 100% | 100% | 3.31% |  |

## Acknowledgements

This work was supported in part by an NIH program to advance systems level understanding of developmental biology (via NIH 1R01DE023575-01A1) and an NSF CAREER award to study morphogen gradient readouts (IOS:1239673).

---

**Figure S1.**
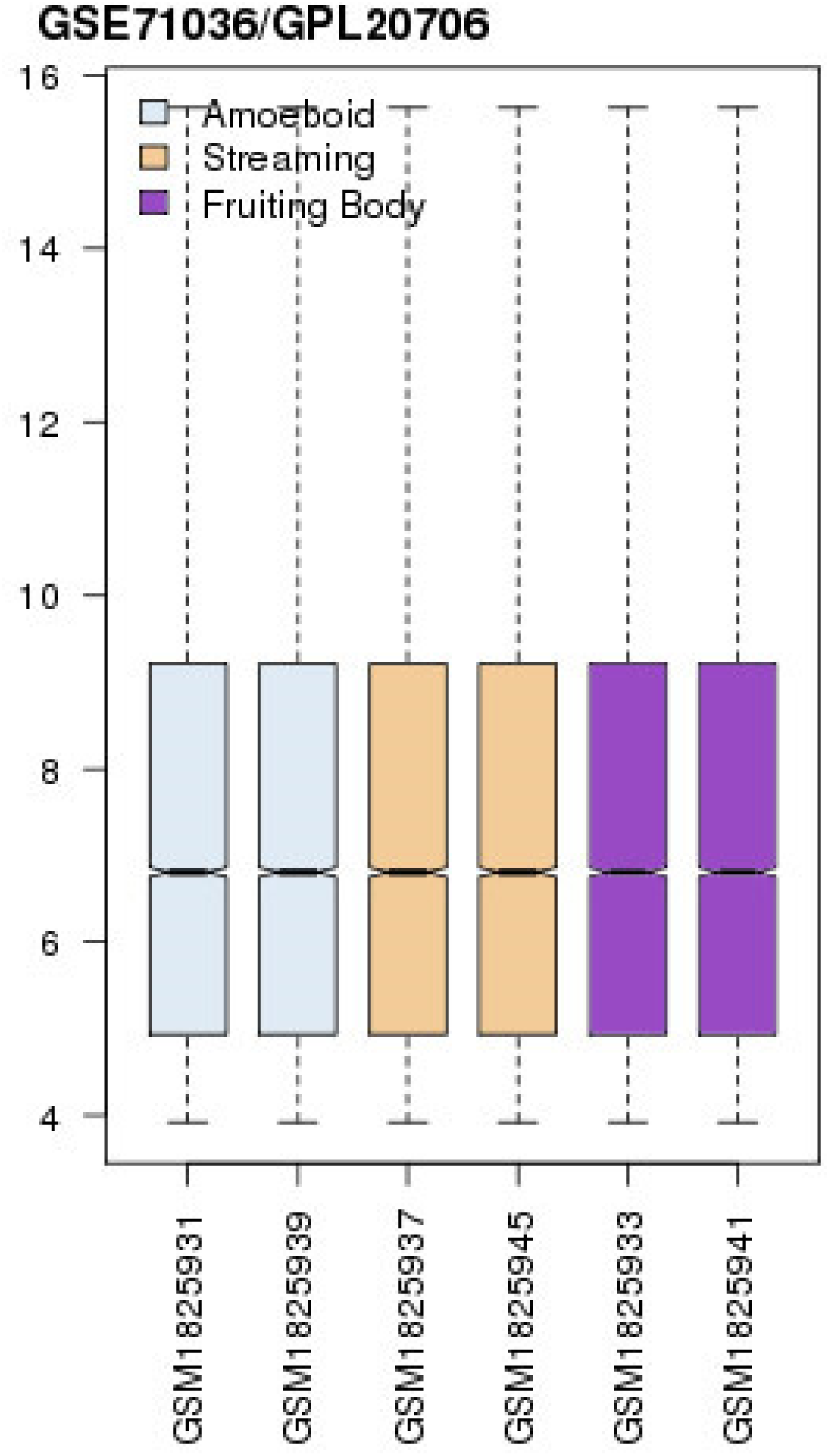
Distributions of expression values for analyzed GSE71036 samples.

**Figure S2.**
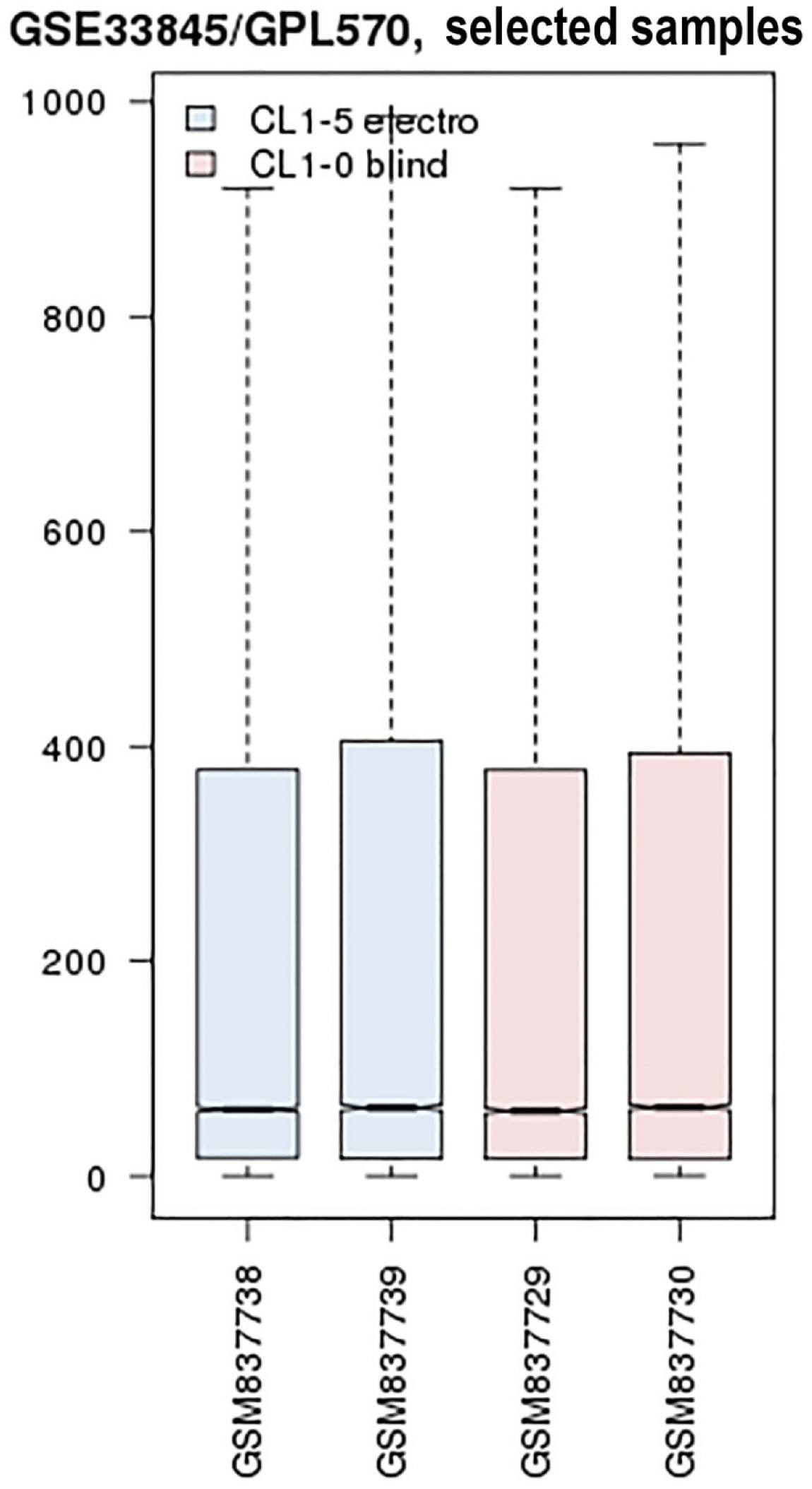
Distributions of expression values for analyzed GSE42407 samples.

## References

Anderson, J. D., 1962 Potassium loss during galvanotaxis of slime mold. J Gen Physiol 45: 567–574.

Barrett, T., D. B. Troup, S. E. Wilhite, P. Ledoux, C. Evangelista et al., 2011 NCBI GEO: archive for functional genomics data sets--10 years on. Nucleic Acids Res 39: D1005–1010.

Barrett, T., S. E. Wilhite, P. Ledoux, C. Evangelista, I. F. Kim et al., 2013 NCBI GEO: archive for functional genomics data sets--update. Nucleic Acids Res 41: D991–995.

Chen, C. Y., Y. H. Jan, Y. H. Juan, C. J. Yang, M. S. Huang et al., 2013 Fucosyltransferase 8 as a functional regulator of nonsmall cell lung cancer. Proc Natl Acad Sci U S A 110: 630–635.

Choi, J. H., H. Y. Jung, H. S. Kim and H. G. Cho, 2000 PhyloDraw: a phylogenetic tree drawing system. Bioinformatics 16: 1056–1058.

Coates, J. C., and A. J. Harwood, 2001 Cell-cell adhesion and signal transduction during Dictyostelium development. J Cell Sci 114: 4349–4358.

Gao, R., S. Zhao, X. Jiang, Y. Sun, S. Zhao et al., 2015 A large-scale screen reveals genes that mediate electrotaxis in Dictyostelium discoideum. Sci Signal 8: ra50.

Haider, S., B. Ballester, D. Smedley, J. Zhang, P. Rice et al., 2009 BioMart Central Portal-unified access to biological data. Nucleic Acids Res 37: W23–27.

Jaffe, L. F., and R. Nuccitelli, 1977 Electrical controls of development. Annu Rev Biophys Bioeng 6: 445–476.

Jones, P., D. Binns, C. McMenamin, C. McAnulla and S. Hunter, 2011 The InterPro BioMart: federated query and web service access to the InterPro Resource. Database (Oxford) 2011: bar033.

Malinovska, L., S. Palm, K. Gibson, J. M. Verbavatz and S. Alberti, 2015 Dictyostelium discoideum has a highly Q/N-rich proteome and shows an unusual resilience to protein aggregation. Proc Natl Acad Sci U S A 112: E2620–2629.

McCaig, C. D., A. M. Rajnicek, B. Song and M. Zhao, 2005 Controlling cell behavior electrically: current views and future potential. Physiol Rev 85: 943–978.

Myre, M. A., A. L. Lumsden, M. N. Thompson, W. Wasco, M. E. MacDonald et al., 2011 Deficiency of huntingtin has pleiotropic effects in the social amoeba Dictyostelium discoideum. PLoS Genet 7: e1002052.

Nakajima, K., K. Zhu, Y. H. Sun, B. Hegyi, Q. Zeng et al., 2015 KCNJ15/Kir4.2 couples with polyamines to sense weak extracellular electric fields in galvanotaxis. Nat Commun 6: 8532.

Nuccitelli, R., M. M. Poo and L. F. Jaffe, 1977 Relations between ameboid movement and membrane-controlled electrical currents. J Gen Physiol 69: 743–763.

Peters, T. W., and M. Huang, 2007 Protein aggregation and polyasparagine-mediated cellular toxicity in Saccharomyces cerevisiae. Prion 1: 144–153.

Reid, B., B. Song and M. Zhao, 2009 Electric currents in Xenopus tadpole tail regeneration. Dev Biol 335: 198–207.

Rice, C., D. Beekman, L. Liu and A. Erives, 2015 The Nature, Extent, and Consequences of Genetic Variation in the opa Repeats of Notch in Drosophila. G3 (Bethesda) 5: 2405–2419.

Saltukoglu, D., J. Grunewald, N. Strohmeyer, R. Bensch, M. H. Ulbrich et al., 2015 Spontaneous and electric field-controlled front-rear polarization of human keratinocytes. Mol Biol Cell 26: 4373–4386.

Shanley, L. J., P. Walczysko, M. Bain, D. J. MacEwan and M. Zhao, 2006 Influx of extracellular Ca2+ is necessary for electrotaxis in Dictyostelium. J Cell Sci 119: 4741–4748.

Smedley, D., S. Haider, S. Durinck, L. Pandini, P. Provero et al., 2015 The BioMart community portal: an innovative alternative to large, centralized data repositories. Nucleic Acids Res 43: W589–598.

Tsai, H. f., C. W. Huang, H. F. Chang, J. J. Chen, C. H. Lee et al., 2013 Evaluation of EGFR and RTK signaling in the electrotaxis of lung adenocarcinoma cells under direct-current electric field stimulation. PLoS One 8: e73418.

Wessels, D., D. F. Lusche, A. Scherer, S. Kuhl, M. A. Myre et al., 2014 Huntingtin regulates Ca(2+) chemotaxis and K(+)-facilitated cAMP chemotaxis, in conjunction with the monovalent cation/H(+) exchanger Nhe1, in a model developmental system: insights into its possible role in Huntingtons disease. Dev Biol 394: 24–38.

Zhang, G., M. Edmundson, V. Telezhkin, Y. Gu, X. Wei et al., 2015 The Role of Kv1.2 Channel in Electrotaxis Cell Migration. J Cell Physiol.

Zhao, m., T. Jin, C. D. McCaig, J. V. Forrester and P. N. Devreotes, 2002 Genetic analysis of the role of G protein-coupled receptor signaling in electrotaxis. J Cell Biol 157: 921–927.

